# Individual Astrocyte Morphology in the Collagenous Lamina Cribrosa Revealed by Multicolor DiOlistic Labeling

**DOI:** 10.1101/2022.12.24.520184

**Authors:** Susannah Waxman, Marissa Quinn, Cara Donahue, Louis D. Falo, Daniel Sun, Tatjana C. Jakobs, Ian A. Sigal

## Abstract

Astrocytes in the lamina region of the optic nerve head play vital roles in supporting retinal ganglion cell axon health. In glaucoma, these astrocytes are implicated as early responders to stressors, undergoing characteristic changes in cell function as well as cell morphology. Much of what is currently known about individual lamina astrocyte morphology has been learned from rodent models which lack a defining feature of the human optic nerve head, the collagenous lamina cribrosa (LC). Current methods available for evaluation of collagenous LC astrocyte morphology have significant shortcomings. We aimed to evaluate Multicolor DiOlistic labeling (MuDi) as an approach to reveal individual astrocyte morphologies across the collagenous LC.

Gold microcarriers were coated with all combinations of three fluorescent cell membrane dyes, DiI, DiD, and DiO, for a total of seven dye combinations. Microcarriers were delivered to 150μm-thick coronal vibratome slices through the LC of pig, sheep, goat, and monkey eyes via MuDi. Labeled tissues were imaged with confocal and second harmonic generation microscopy to visualize dyed cells and LC collagenous beams, respectively. GFAP labeling of DiOlistically-labeled cells with astrocyte morphologies was used to investigate cell identity. 3D models of astrocytes were created from confocal image stacks for quantification of morphological features.

DiOlistic labeling revealed fine details of LC astrocyte morphologies including somas, primary branches, higher-order branches, and end-feet. Labeled cells with astrocyte morphologies were GFAP^+^. Astrocytes were visible across seven distinct color channels, allowing high labeling density while still distinguishing individual cells from their neighbors. MuDi was capable of revealing tens to hundreds of collagenous LC astrocytes, in situ, with a single application. 3D astrocyte models allowed automated quantification of morphological features including branch number, length, thickness, hierarchy, and straightness as well as Sholl analysis.

MuDi labeling provides an opportunity to investigate morphologies of collagenous LC astrocytes, providing both qualitative and quantitative detail, in healthy tissues. This approach may open doors for research of glaucoma, where astrocyte morphological alterations are thought to coincide with key functional changes related to disease progression.

## 1. Introduction

### 1.1. Background

The lamina cribrosa (LC) is an early site of retinal ganglion cell (RGC) injury in glaucoma, a leading cause of irreversible blindness worldwide. The LC is densely populated by astrocytes which, in health, play vital roles in maintaining RGC homeostasis and healthy vision(J. Salazar et al., 2019; Prada et al., 2016). Growing evidence supports an early role of lamina astrocyte functional and morphological changes as factors that can be protective against and can contribute to glaucomatous neurodegeneration.(Hernandez et al., 2008; Liu et al., 2022; Morgan, 2000; Shinozaki and Koizumi, 2021; Sun et al., 2017) Changes in astrocyte function are known to impact several key processes involved in optic nerve head (ONH) health, such as hemodynamic regulation,(Attwell et al., 2010; Dong et al., 2022) blood-brain barrier maintenance,(Heithoff et al., 2021; Liu et al., 2018) and extracellular matrix remodeling.(Hernandez, 2000; Schneider and Fuchshofer, 2016) These astrocyte functional changes are associated with changes in cell morphology. Astrocytes undergo extensive reorganization in the transition between health and glaucoma. Characteristic “reactive” lamina astrocytes exhibit thickened and shortened cell processes, loss of higher-order branches, and reduced spatial coverage of individual astrocytes(Lye-Barthel et al., 2013; Sun et al., 2010; Sun and Jakobs, 2012; Wang et al., 2017). The morphology and arrangement of astrocytes across the lamina can provide important insights into their roles in health and in disease progression.

### 1.2. Why is a new method to label astrocytes needed?

Established methods to analyze LC astrocyte morphology have important limitations. Despite the recognized importance of LC astrocytes in health and glaucoma, the methods used to investigate their morphology have critical constraints that inhibit effective large-scale analysis of individual cells. Mouse lines expressing fluorescent reporters under the control of astrocyte-specific promoters(Lye-Barthel et al., 2013; Nguyen et al., 2017; Wang et al., 2017) allow for high-quality visualization of these cells. However, mice lack a true collagenous LC like that found in human eyes. Collagen beams and neural tissue pores of the collagenous LC define crucial biological and mechanical properties of the tissue(May and Lütjen-Drecoll, 2002; Midgett et al., 2020; Oikawa et al., 2021; Sigal and Ethier, 2009) that the analogous structure in mice, the glial lamina, does not approximate. To visualize astrocytes of the collagenous LC, the research model must have a true collagenous LC.

A common method used to visualize and identify astrocytes in LC tissue sections is immunohistochemistry of the GFAP protein. GFAP expression of astrocytes in the brain tends to be low in health and is upregulated in reactivity.(Liddelow and Barres, 2017) In the glial lamina, GFAP is strongly expressed by astrocytes, even in the absence of damage or reactive gliosis.(Lye-Barthel et al., 2013; Sun et al., 2009) Immunohistochemical labeling of GFAP reveals the location of the GFAP^+^ intermediate filaments found in most astrocytes. However, when using methods that label all or most astrocytes in the LC, it is often not possible to distinguish one cell from its neighbor due to the high density of astrocytes in the region and their complex and branched organizations. Complexity is further compounded by the fact that GFAP labeling does not reveal the full extent of astrocyte morphology, as has been shown in the brain, retina, and mouse glial lamina.(Bushong et al., 2002; Escartin et al., 2021; Luna et al., 2016; Stokum et al., 2018; Sun and Jakobs, 2012) GFAP can be visualized in many thicker primary processes, but often does not label finer higher-order processes, missing important morphological detail. Astrocyte GFAP content and organization changes substantially in glaucoma,(Chaudhary et al., 2022; Ling et al., 2020) but reveals only a fraction of astrocyte morphology. Serial block-face scanning electron microscopy can produce high enough resolution images for users to reconstruct the entirety of individual astrocytes and separate them from their neighbors with excellent detail.(Bonney et al., 2022; Calì et al., 2019) However, understanding the diversity of astrocyte morphologies and spatial relationships to other structures across the LC of several individuals with this approach would be prohibitively time- and resource-intensive.

In lieu of large animal astrocyte reporter lines, astrocyte gene transfer approaches(Ge et al., 2020; Merienne et al., 2013) could be attempted in the collagenous LC of research animals such as non-human primates, *in vivo*. This would require considerable time, cost, and risk. The LC can be difficult to target in live animals both due to selectivity of the blood-brain barrier and the poorly accessible location of the structure in the eye. Single-cell dye injections can bypass many challenges of gene transfer approaches and similarly reveal astrocyte morphology. This approach has been successfully implemented in astrocytes of the brain(Bushong et al., 2002; Moye et al., 2019) and the mouse glial lamina(Butt et al., 1994; Sun, 2018) specifically. However, as this technique requires cannulating cells with a micropipette one at a time, it can be time and labor-intensive to visualize multiple cells across the tissue at once.

### 1.3. How does Multicolor DiOlistic labeling fit in as a logical next step?

DiOlistic labeling is a technique that utilizes a gene gun to propel gold or tungsten microcarriers coated in cell membrane dyes into individual cells within tissues. With this approach, tens to hundreds of dye-coated microcarriers can be delivered to tissues simultaneously. This technique has been implemented in the brain and retina to visualize neuron morphologies(O’Brien and Lummis, 2006; Santina and Ou, 2018; Seabold et al., 2010; Staffend and Meisel, 2011) and in the mouse glial lamina(Jakobs, 2014; Sun et al., 2010) to visualize astrocyte morphologies. By delivering microcarriers coated in all combinations of 3 spectrally distinct fluorescent cell membrane dyes (DiI, DiD, and DiO), a multicolor DiOlistic (MuDi) labeling approach has been accomplished in the mouse brain and retina.(Gan et al., 2000) These multiple separate color channels enabled labeling and visualization of a large number of cells in close proximity to each other while still allowing to distinguish cells from their neighbors.

In this work, we aimed to determine the suitability of MuDi labeling(Gan et al., 2000) as an approach to visualize collagenous LC astrocyte morphology, *in situ*. The scope of this work is focused on testing the ability of MuDi labeling to reveal individual astrocyte morphologies in the collagenous LC. We demonstrate that this technique reveals morphological detail of individual astrocytes across the LC with a single labeling application. Astrocyte morphological features can be semi-automatically quantified and extracted from images on a cell-by-cell and branch-by-branch basis for detailed analysis. This labeling method rendered samples compatible with other labeling and imaging modalities such as immunolabeling for GFAP, lectin-labeling of blood vessels, and second harmonic generation (SHG) imaging of LC collagen beams. The biological relevance of findings from LC astrocyte models will be discussed in detail in future work. MuDi labeling of the collagenous LC can open new avenues for qualitative and quantitative investigation of astrocyte morphology in health, and in the future, in glaucoma.

## 2. Methods

### 2.1. Tissue acquisition and preparation

Pig, sheep, goat, and monkey eyes were utilized for experiments (N = 2, 4, 16, and 12 eyes, respectively). Pig, sheep, and goat eyes were acquired within 1 hour of sacrifice. Rhesus macaque monkey eyes were obtained within 30 minutes of sacrifice. ONHs were isolated and fixed in methanol-free 4% PFA for 2-24 hours and held in PBS at 4°C. All procedures were approved by the University of Pittsburgh’s Institutional Animal Care and Use Committee (IACUC) and adhered to both the guidelines set forth in the National Institute of Health’s Guide for the Care and Use of Laboratory Animals and the Association of Research in Vision and Ophthalmology (ARVO) statement for the use of animals in ophthalmic and vision research. No live animals were used in this work.

### 2.2. Vibratome sectioning

ONHs were embedded in low melting temperature agarose (Sigma, A9045), secured to the stage of a vibratome (Leica, VT1200S) with cyanoacrylate glue, and immersed in ice-cold PBS. Coronal slices were cut at 150μm-thickness. As the collagen of the ONH is highly elastic, high blade advancement speeds can cause deformation of the tissue before cutting, resulting in uneven slices, increased disruption of cell membranes, and in some instances, dislodging of the tissue from its agarose block, resulting in loss of tissue orientation. Low blade advancement speed (0.01-0.05 mm/second) helped ensure even tissue cutting. Slices were transferred from the vibratome, gently with a paintbrush, to fresh PBS. Slices were screened under polarized light and brightfield microscopy to visualize the collagen beams and surrounding anatomical structures indicative of the LC region. Slices were stored in PBS at 4°C for up to 48 hours prior to DiOlistic labeling.

### 2.3. Multicolor DiOlistics bullet preparation

DiOlistics bullets were prepared similarly to as done previously by others.(Benediktsson et al., 2005; Gan et al., 2000; Jakobs, 2014; O’Brien and Lummis, 2006; Sun et al., 2010) For multicolor bullets, 70mg of 1.0μm diameter gold microparticles (BioRad, 1652263) were divided equally into 7 1.5mL tubes and each mixed in 100μL ethanol. 4mg of each DiI (Sigma-Aldrich 42364, 549/567nm), DiO (Invitrogen D275, 484/501nm), and DiD (Invitrogen D7757, 644/665nm excitation/emission) were used to make separate stock solutions in 500μL ethanol. Dye solutions were added to tubes containing gold microparticles in all combinations and in equal proportions, such that each tube received a total of 1.7mg of dye. For example, DiI-coated microparticles received 1.7mg of DiI. DiI + DiO-coated microparticles received 0.85mg of DiI and 0.85mg of DiO. DiI + DiO + DiD-coated microcarriers received 0.57mg of each dye.

Gold-dye mixtures were vortexed for 30 seconds, pipetted onto separate glass microscope slides (Fisher, 12550A3), and spread evenly across slides with a pipette tip. Ethanol was allowed to fully evaporate for at least 20 minutes, rendering dry microcarriers coated in dyes. The dry microcarriers were scraped off their respective slides with a razor and collected together. Microcarriers were suspended in 1mL deionized water and sonicated for 10 minutes to minimize clumping. Sonicated microcarriers were filtered through a 20μm filter (Millipore, NY2004700) to remove large clumps and resuspended in 3mL deionized water. 30μL of 0.5 mg/ml polyvinylpyrrolidone in isopropanol was added to this mixture.

Microcarrier mixture was loaded into Tefzel tubing (Bio-Rad, 1652441) in a tubing prep station (Bio-Rad, 1652418). Microcarriers were allowed to sediment to the bottom of the tube for 5-10 minutes before liquid was slowly withdrawn from the tube. Tubing was rotated for 5 minutes in the tubing prep station to distribute the microcarriers along the interior of the tubing. Nitrogen was run through the tubing for 30 minutes at 2 PSI to remove the remaining water. Tubing was cut into 1.3cm lengths with a tubing cutter (Bio-Rad, 1652422) and stored in an airtight container with desiccant, protected from light, for as long as 6 months. Single-color bullets were prepared the same way as multicolor bullets, but using 4mg of DiI or DiD as the only dye and 23mg of gold microparticles instead of 70mg.

### 2.4. DiOlistic labeling

A Helios Gene Gun (BioRad, 1652411) was loaded with bullets, connected to a pressurized nitrogen tank, and used to deliver dye-coated microcarriers into vibratome slices at 150-200 PSI. Vibratome slices were gently removed from PBS with a paintbrush and placed in a 60mm dish. Slices were covered with a 20μm filter (Millipore, NY2004700) and the spacer of the gene gun was positioned flush with the filter. The gene gun was oriented perpendicular to the sample as in Figure 2, and each sample was shot once with the gene gun. Samples were returned to PBS within 30 seconds to prevent sample dehydration and tissue damage. Samples counterstained with DAPI were incubated in 1:1000 DAPI (ThermoFisher, 62248) for 20 minutes and washed 3 times with PBS. Slices were mounted labeled-side-up on glass slides in water-based mounting medium (Shandon, 9990402) for microscopy.

### 2.5. Additional non-DiOlistic labeling

For samples labeled immunohistochemically and/or with lectin (non-DiOlistic labeling), this was done either before, after, or without DiOlistic labeling. Each method has specific benefits and limitations, detailed below.

#### Before DiOlistic labeling

Samples with non-DiOlistic labeling completed before DiOlistic labeling, such as those in **Figure 1C**-**E**, allowed imaging of DiOlistically-labeled cells and other labels in the same imaging session. This minimizes the risk of interpretation errors, such as false negative and false positive colocalization of GFAP and DiOlistics signals, caused by potential image registration imperfections or sample distortion between imaging sessions. These samples however, cannot be extensively permeabilized with detergents to enhance non-DiOlistic label penetration through sample thicknesses. Detergents, such as the commonly used Triton X-100, create holes in cell membranes that can result in substantial leakage of DiOlistically-delivered dyes from cells into extracellular spaces and onto neighboring cells, preventing clear visualization of individual cell morphologies. This method is also not readily compatible with the use of multicolor DiOlistic bullets, as many fluorescent probes used in immunohistochemistry have similar fluorescent profiles to DiD, DiO, or DiI. Therefore, single-color bullets were used in these samples and less cells were labeled at once to ensure clear spatial separation of labeled cells.

**Figure 1.**
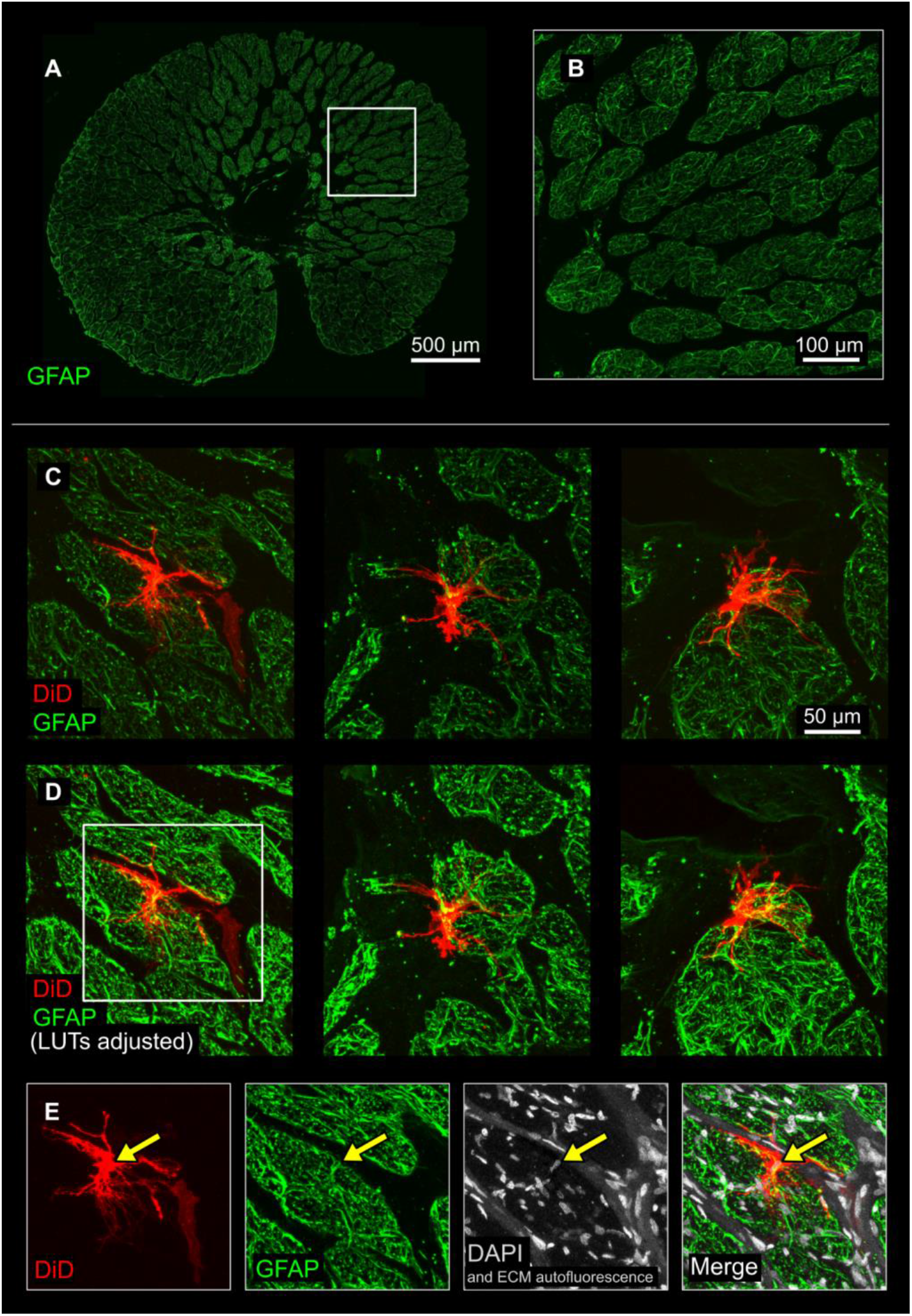
DiOlistically-labeled cells with astrocyte morphologies were GFAP^+^. **A**) GFAP-immunolabeled vibratome slice through the ONH. Astrocytes and their GFAP^+^ intermediate filaments densely populate neural tissue pores and line extracellular matrix beams in the LC. **B**) Detail of the white box in **A** shows the complex, filamentous, and tightly packed arrangement of GFAP filaments within the LC. IHC of GFAP reveals a limited amount of astrocyte morphology, labeling only a portion of the cell. This, coupled with the complexity, scale, and density of GFAP^+^ filaments renders questions about individual astrocyte morphologies difficult to investigate. **C**) Fluorescent cell membrane tracers, such as DiD, can be introduced into individual cells through single-color DiOlistic labeling. This revealed the morphology of individual GFAP^+^ astrocytes within the context of surrounding tissues. **D**) Images from C shown with LUTs adjusted to improve visibility of GFAP colocalization in regions with strong DiOlistics signal and of fine astrocyte branches with less intense DiOlistics signal. **E**) Astrocyte within the white box in **D** with DiD, GFAP, and DAPI channels shown separately and merged. Yellow arrows indicate location of the DiOlistically-labeled cell’s nucleus. Representative images were from goat LC. Images are maximum intensity projections of confocal Z stacks. **A**: 20x, **B-E**: 40x.

Three vibratome slices from three goat eyes were incubated for 4 hours in 5% BSA blocking solution without permeabilization agent, overnight in GFAP primary antibody (rabbit origin, (Invitrogen, PA5-16291) at 1:200 in blocking solution, and washed 3×10min in PBS. Samples were incubated with secondary antibody at 1:500 dilution (488 goat-anti-rabbit, Abcam, ab150077) for 6 hours at room temperature, washed 3×10min in PBS, incubated in 1:1000 DAPI for 20 minutes, and washed 3×10 minutes in PBS. Immediately after non-DiOlistic labeling was completed, samples were labeled with single-color DiD DiOlistics.

#### After DiOlistic labeling

Samples labeled with non-DiOlistic labels after DiOlistic labeling, such as those in **Figure 8**, were permeabilized in order to enhance non-DiOlistic label penetration, without the same concern necessary for cell membrane integrity as samples treated with non-DiOlistic labels before DiOlistic labeling. Samples treated with non-DiOlistic labels after DiOlistic labeling must however be removed from their slides after imaging of DiOlistics signal for this additional labeling and washing. Images from the first round of imaging (to visualize DiOlistic labeling) and second round of imaging (to visualize non-DiOlistic labeling) must be carefully registered to each other based on shared information between the two, such as DAPI signal and/or ECM autofluorescence. The process of lipophilic dye removal and re-labeling caused minimal tissue distortion but given the scale of astrocytes and their fine processes, interpretation of colocalized signals should be done with caution and/or verified with samples labeled before to DiOlistics.

After imaging of 4 DiOlistically-labeled slices was completed, one slice was introduced to an ethanol gradient (50%, 70%, 90% for 1 hour each) and held in 98% ethanol for two additional days to test the ability of ethanol to remove dyes from the tissue. When this sample was observed post-ethanol wash with the same imaging parameters, no DiOlistically-labeled cells or appreciable MuDi dye signal was observed in the tissue. The remaining three samples were blocked, labeled with non-DiOlistics labels, and washed in PBS prior to ethanol-mediated MuDi dye removal. These three samples were blocked and permeabilized thoroughly in 5% BSA 0.05% Triton X-100 (block/perm) for 24 hours, incubated with GFAP primary antibody diluted 1:200 and 1:200 DyLight Tomato lectin (VectorLabs, DL-1178) in block/perm for 48 hours, washed for 1 day in PBS, incubated for 24 hours in 1:500 secondary in block/perm, and washed 3×10 minutes in PBS. Samples were then introduced to the same ethanol gradient as above. Samples were rehydrated for imaging and mounted in water-based mounting medium with the previously imaged side facing up.

#### Without DiOlistic labeling

Tissue labeled for GFAP and without DiOlistic labeling, to visualize GFAP alone (**Figure 1A, B**, N = 1 eye), was prepared the same way as samples labeled after DiOlistic labeling, but without lectin or ethanol washes.

### 2.6. Imaging

An upright confocal microscope (BX61; Olympus) was used to screen each DiOlistically-labeled sample at low magnification (4x, 10x, and/or 20x). In samples with sufficient density and quality of labeling in screening, tissues were observed under high magnification (40x). Dyes must distribute across the cell membrane of individual cells in order to reveal morphology. Cells were screened at 40x magnification for extent of dye spread prior to extensive imaging. Segmentation-quality images were collected after astrocyte end-feet and high-order processes could be visualized across the LC. This occurred within 1 hour of labeling at room temperature. Imaging was completed within 24-48 hours of DiOlistic labeling. 40x images were collected at an XY image size of 1024×1024 or 800×800 pixels, and with a 0.5-2μm Z step throughout the visible thickness of the sample. Imaging of non-DiOlistics labels (GFAP, lectin) was completed with the same confocal microscope at 20 and 40x magnifications, with the same X, Y, and Z image sizes as used for imaging of DiOlistics. SHG imaging was conducted with a multiphoton microscope (A1; Nikon) at 25x. SHG images were collected at a 1024×1024 or 512 × 512 pixel XY image size at a 2-5μm Z step throughout the visible thickness of the sample.

### 2.7. Astrocyte segmentation and 3D model construction

Images were imported into Imaris (Bitplane, version 9.9) software for 3D visualization and model construction of labeled astrocytes. The Filaments tool and the “AutoPath-no loops” algorithm were used to automatically segment individual astrocytes. Each astrocyte was created with a single starting-point at its soma and seed-points along its branches. Imaris-generated models were checked to ensure that they faithfully represented image volume information. Aberrant branches were removed, and missing branches were added manually. Quantitative branch-by-branch level information about morphological features, such as branch number, length, thickness, hierarchy, straightness, and Sholl intersections was extracted from the Statistics panel. Sholl analysis was conducted with the Filament Sholl Analysis XTension.

## 3. Results

### 3.1. DiOlistically-labeled cells with astrocyte morphologies are GFAP^+^

GFAP labeling revealed astrocyte intermediate filaments in the collagenous LC. However, GFAP labeling was insufficient to understand the morphology of individual astrocytes within LC tissues (**Figure 1A, B**). The high density and structural complexity of GFAP filaments rendered discerning one astrocyte from its neighbor not possible. Additionally, GFAP does not reveal the full extent of the spatial territory occupied by astrocytes.(Bushong et al., 2002; Escartin et al., 2021; Luna et al., 2016; Stokum et al., 2018; Sun and Jakobs, 2012) GFAP filaments are often found within thicker primary processes but can be absent in higher-order processes. Conclusions about the morphology of astrocytes labeled with GFAP alone are therefore subject to critical misinterpretations.

DiOlistic delivery of dye-coated microcarriers to GFAP-labeled goat LC tissues resulted in stochastic cell membrane labeling of individual cells. The density of single-color dye-coated microcarriers delivered to LC tissues was optimized to ensure clear spatial separation of labeled cells. DiOlistically-labeled cells with astrocyte morphologies colocalized partially with GFAP, demonstrating astrocyte identity. **Figure 1C & D** show instances of DiOlistically-labeled cells of the goat LC with astrocyte morphologies. These cells had a single soma, typically 3-8 major processes, and multiple fine higher-order processes. Processes extended within neural tissue pores and contacted LC beams. In samples counterstained with DAPI, these cells contained a single nucleus (**Figure 1E**).

### 3.2. Multicolor DiOlistic labeling of the LC: workflow

To visualize multiple collagenous LC astrocytes in close proximity to each other while still discerning one cell from another, we employed a MuDi labeling approach, as done by Gan et al(Gan et al., 2000) to visualize cells in the brain and retina of mice. ONHs from pig, sheep, goat, and monkey eyes were sectioned into 150 μm-thick coronal slices with a vibratome (**Figure 2A**). Groups of microcarriers were coated with all combinations of 3 spectrally distinct fluorescent cell membrane dyes to create a total of seven color combinations (**Figure 2B**). Groups of tens to hundreds of multicolor dye-coated microcarriers were loaded into “bullets” for simultaneous delivery to tissues with a gene gun. These groups of microcarriers were propelled into tissue slices containing collagenous LC (**Figure 2C**). Microcarriers were embedded within cells and the dyes they carried were distributed across individual cell membranes. DiOlistically-labeled tissues were imaged with confocal microscopy to visualize labeled cells and DAPI-labeled cell nuclei and/or ECM autofluorescence. Select tissues were imaged with second harmonic generation (SHG) microscopy to visualize LC collagen (**Figure 2D**).

**Figure 2.**
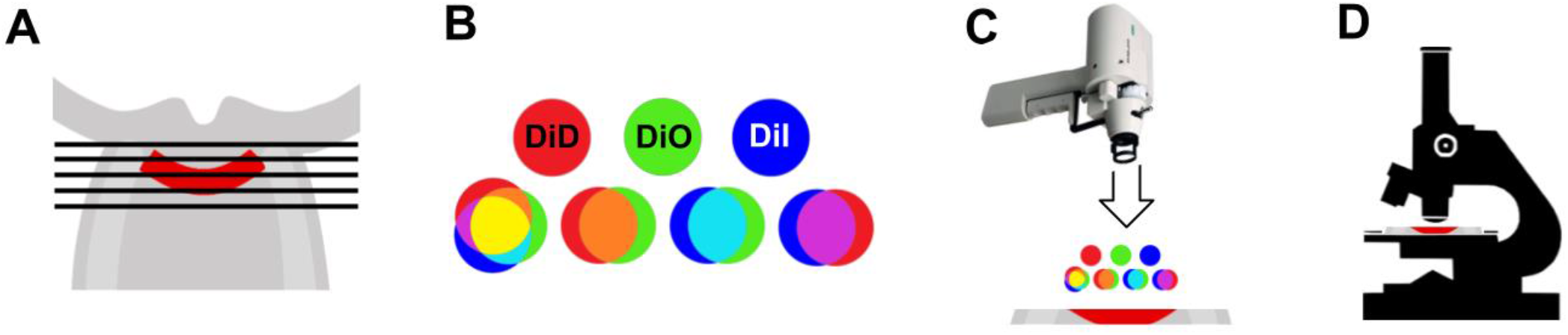
Multicolor DiOlistic labeling of the lamina cribrosa: workflow. **A)** Optic nerve heads from goat, sheep, pig, and monkey eyes were sectioned coronally at 150μm-thickness to collect slices containing LC (red). **B)** Gold microcarriers 1μm in diameter were coated with all combinations of 3 fluorescent cell membrane dyes, DiD, DiO, and DiI, to create a total of 7 distinct color channels. **C)** A gene gun was used to propel groups of dye-coated microcarriers into vibratome slices containing LC. **D)** Labeled tissues were imaged with confocal microscopy, and/or second harmonic generation microscopy.

### 3.3. Multicolor DiOlistic labeling allows LC astrocyte visualization across seven distinct color channels

MuDi allowed for LC astrocyte visualization across the seven distinct color channels created by all combinations of the fluorescent dyes, DiD, DiO, and DiI (**Figure 3**). The seven color channels shown in **Figure 3** were the most commonly observed colors for labeled astrocytes. Example cells in **Figure 3** were of goat origin. As noted in Gan et al,(Gan et al., 2000) additional colors were observed in cells labeled by multiple microcarriers. For example, a cell in this study shown in cyan received one part DiI and one part DiO. If this cyan cell received an additional microcarrier containing one part DiI and one part DiD, it would appear as an intermediate color not included in the original seven categories. At the density of microcarriers included in each bullet, the gas pressure used for each gene gun shot, and the distance the gene gun was set from the tissues in this study, most labeled cells received only one microcarrier. The distribution of microcarriers delivered to tissue samples could be observed under transmitted light (**Supplemental Figure 1**) as dark puncta approximately 1μm in diameter.

**Figure 3.**
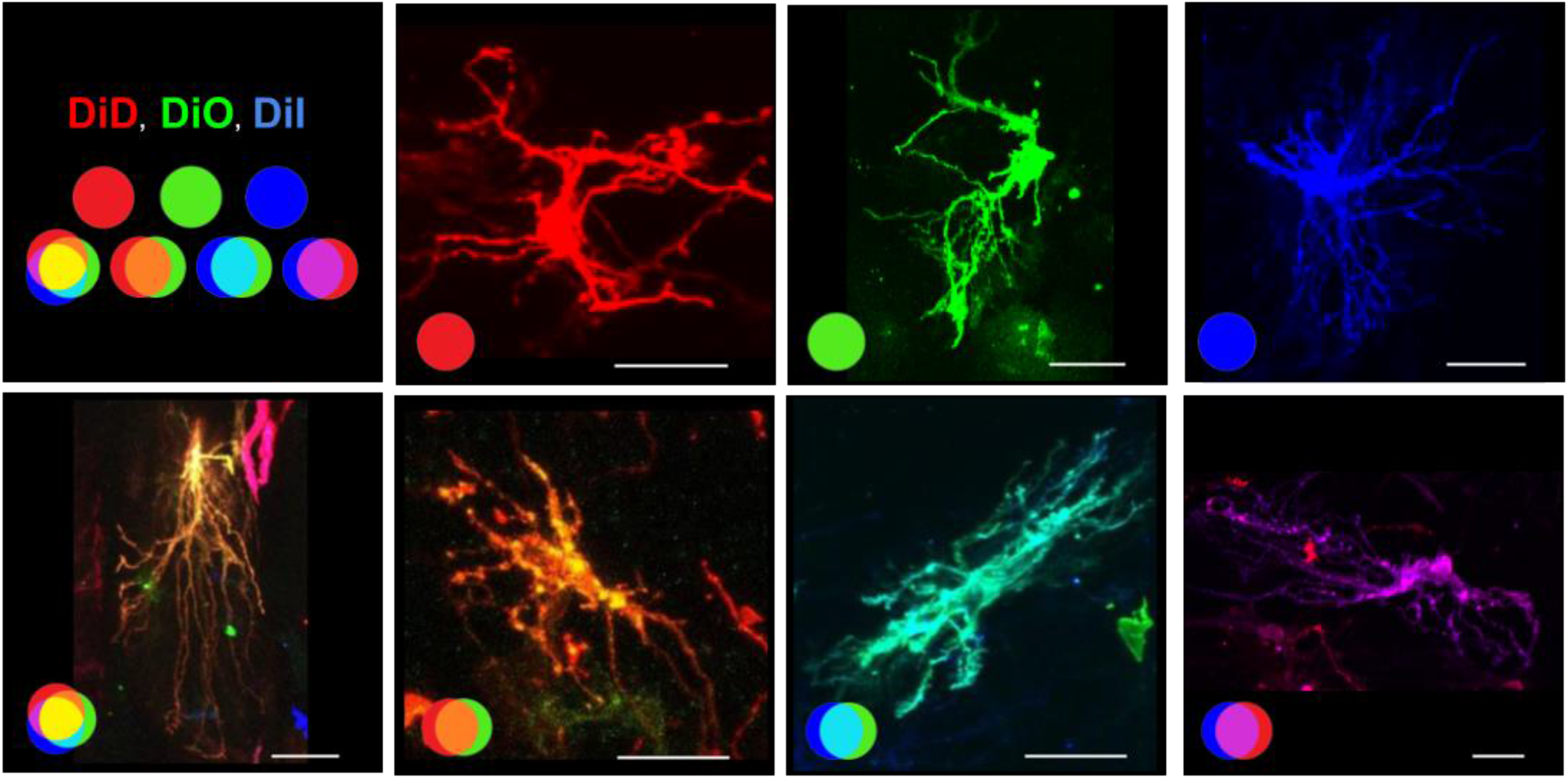
Multicolor DiOlistic labeling allows individual LC astrocyte visualization across seven distinct color channels Individual LC astrocytes of the goat labeled with MuDi at 40x magnification. All combinations of the fluorescent dyes, DiD, DiO, and DiI, allowed cells to be visualized in distinct color channels depending upon the dye(s) they received. Images are maximum intensity projections of confocal stacks. DiD: red, DiO: green, DiI: blue. Scale bar: 25 μm.

### 3.4. Multicolor DiOlistic labeling reveals 3D morphologies of individual astrocytes across the LC

MuDi-labeled coronal slices of the LC allowed for visualization of tens to hundreds of individual astrocytes across the tissue, *in situ* (**Figure 4A**). Confocal images collected at high magnification revealed cell bodies and fine processes of individual cells in close proximity to each other (**Figure 4B**). Most astrocyte branches were oriented along the coronal plane while some spanned the anterior-posterior direction, as seen in mice.(Lye-Barthel et al., 2013; Wang et al., 2017) Example tissue shown in **Figure 4** was of goat origin. Optimal labeling density was approximately 140 cells per mm^2^ of canal section area. At this labeling density, we were able to visualize a large number of cells across the LC with minimal instances of two or more cells labeled with the same dye(s) in close enough proximity as to obscure individual cell morphologies. Collagen beams characteristic of the collagenous LC were visualized with SHG, revealing the relationship of DiOlistically-labeled astrocytes to surrounding collagen supports. SHG signal colocalized with tissue autofluorescence in the 405nm channel observed in confocal microscopy and aided in registration of SHG images to confocal images.

**Figure 4.**
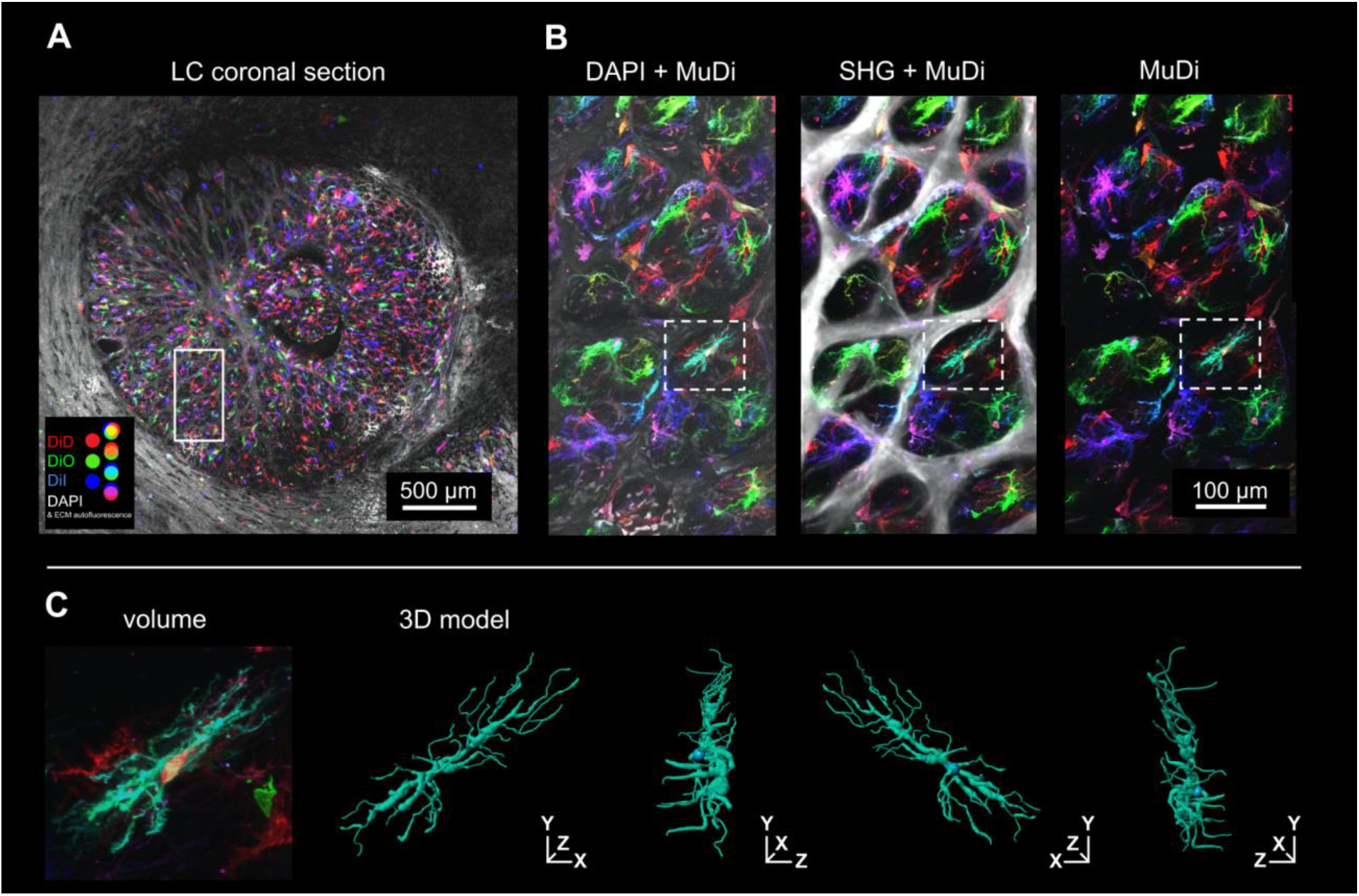
Multicolor DiOlistic labeling reveals 3D morphologies of individual astrocytes across the LC MuDi reveals 3D morphologies of individual astrocytes across the LC. **A**) A coronal section through the goat LC labeled with MuDi. Dye-coated microcarriers labeled hundreds of cells across the sample. Image collected at 10x magnification. **B**) High magnification of the white box within **A** reveals fine details of individual cells within LC pores. DAPI-labeled nuclei and autofluorescence from LC beams (left). Collagen imaged via SHG overlaid verifies the location and appearance of LC beams and neural tissue pores (middle). The MuDi channels are shown alone (right). **C**) Detail of the astrocyte within the white box in **B**. 3D segmentation of the same cell is shown at anterior, sagittal, and posterior viewpoints. Confocal and SHG images are shown as maximum intensity projections. DiD + DiO + DiI = MuDi

Z-stack images collected at 40x magnification allowed for creation of DiOlistically-labeled astrocyte 3D models (**Figure 4C**). Example astrocyte models are shown in relation to their respective confocal volume information in **Figure 5** and **Video 1**. These models granted automated extraction of quantitative morphological features such as branch length, thickness, hierarchy, and straightness (**Figure 6**). Three example astrocyte models from the goat LC are included in **Figure 6**. For 3D views of these astrocytes see **Video 2**. The same methodology was used to create 50 astrocyte models, supporting the robustness of this method for systematic analysis. Sholl analysis (**Figure 7**), convex hull area, number of branching points, number of terminal points, cell volume, and a variety of other quantitative metrics were also readily captured. Sholl data from 3 example collagenous LC astrocytes demonstrates a similar average distribution of crossing counts to mouse glial lamina astrocytes from Lye-Barthel et al, (Lye-Barthel et al., 2013) with higher crossing counts in collagenous LC astrocytes. The ability to extract these quantitative metrics can be helpful in future studies designed to characterize and/or compare healthy and glaucomatous collagenous LC astrocyte morphologies.

**Figure 5.**
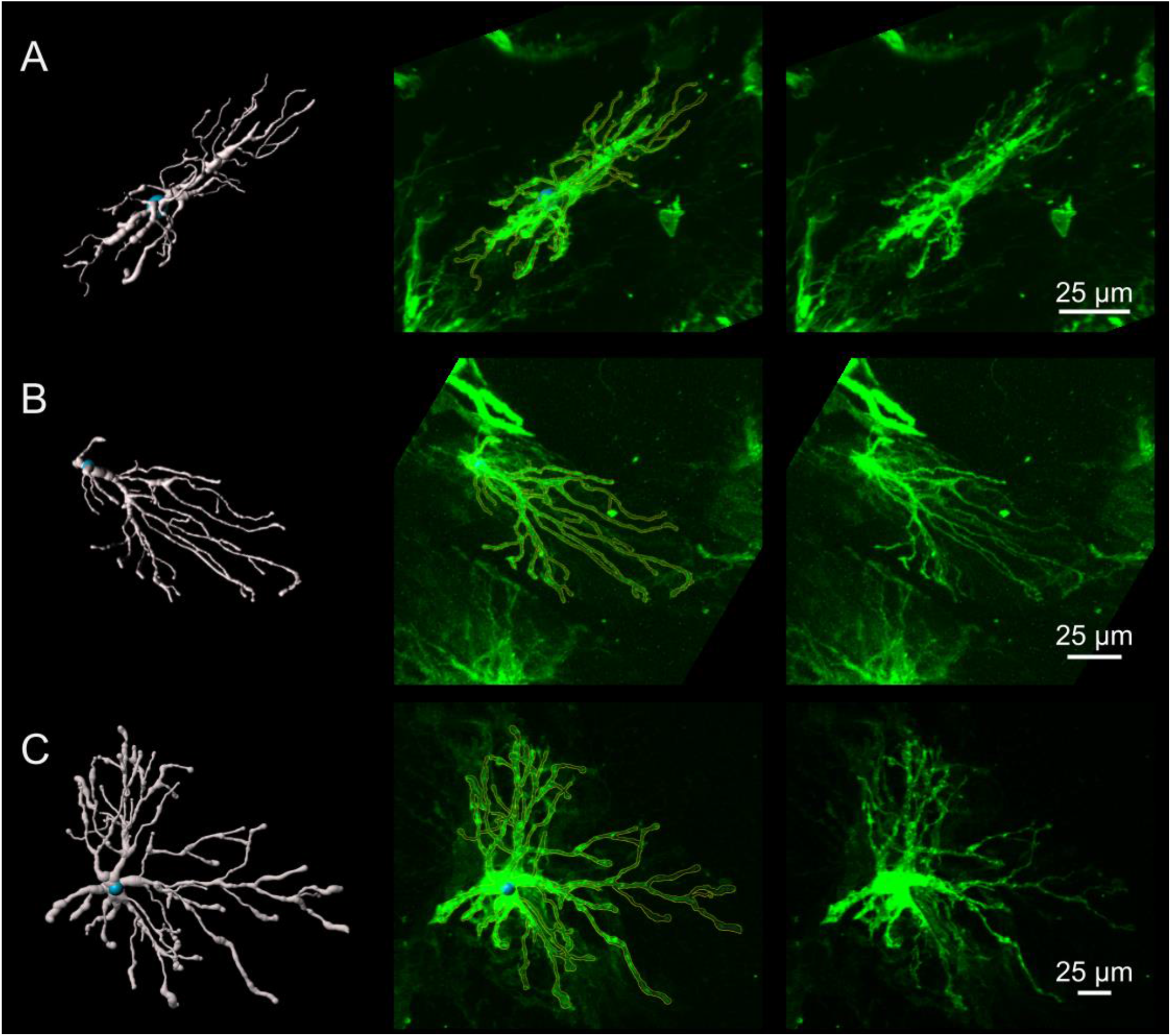
3D astrocyte models reflect respective confocal images. Example goat astrocytes in A-C are shown as 3D models (left), model outlines (yellow) overlaid on their respective confocal images (middle), and as confocal images alone (right). Confocal images here were each pseudocolored green to facilitate uniform visualization.

**Figure 6.**
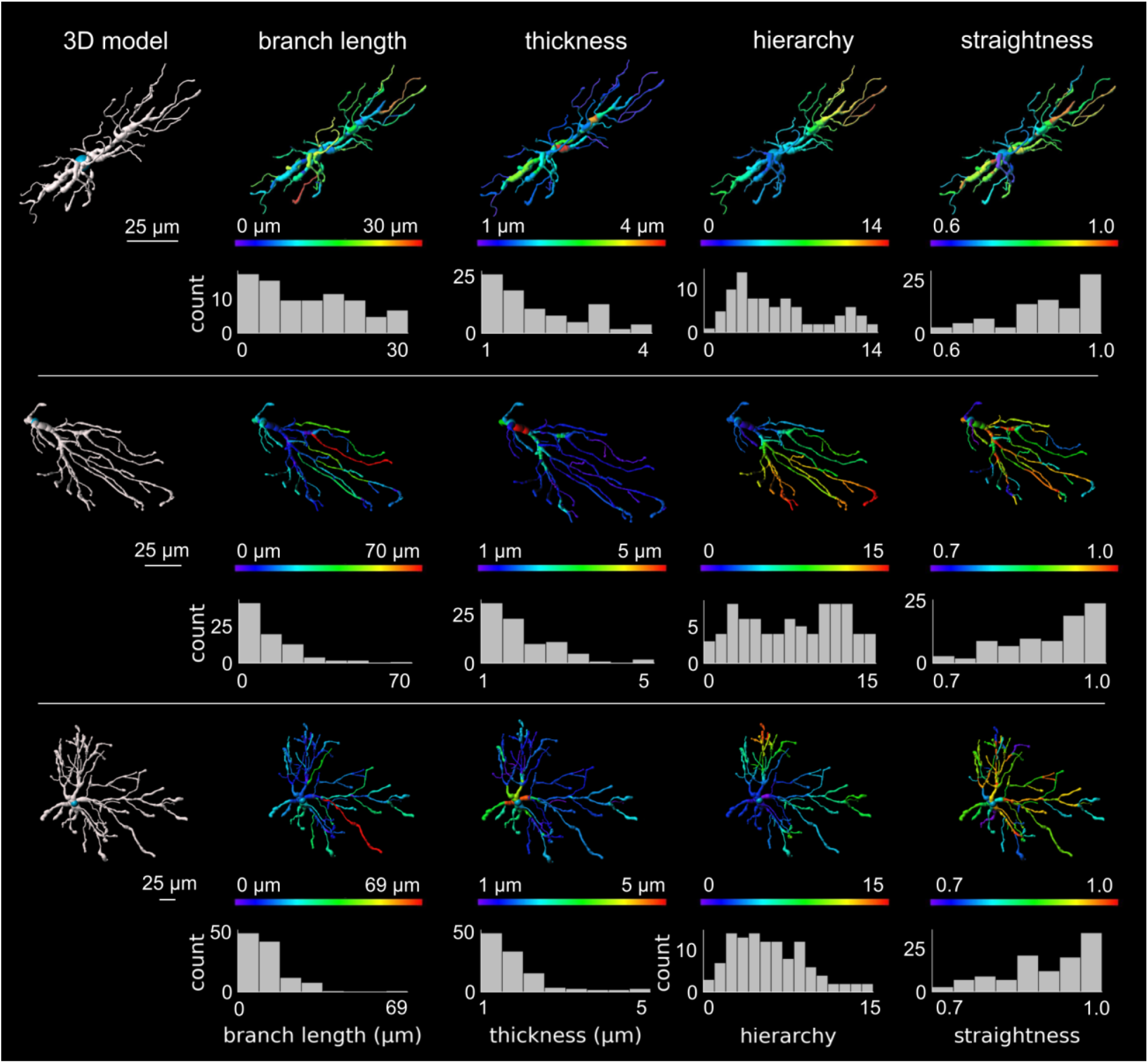
3D astrocyte segmentations allow extraction of morphological features for quantitative analysis. Example astrocyte 3D segmentations (shown in white) were used for automated extraction of morphological features including astrocyte branch length, thickness, hierarchy, and straightness. These morphological features were readily visualized on their respective 3D models (colormaps) and extracted for future statistical analysis at a cell-by-cell and branch-by-branch level (histograms). Example cells were of goat origin.

**Figure 7.**
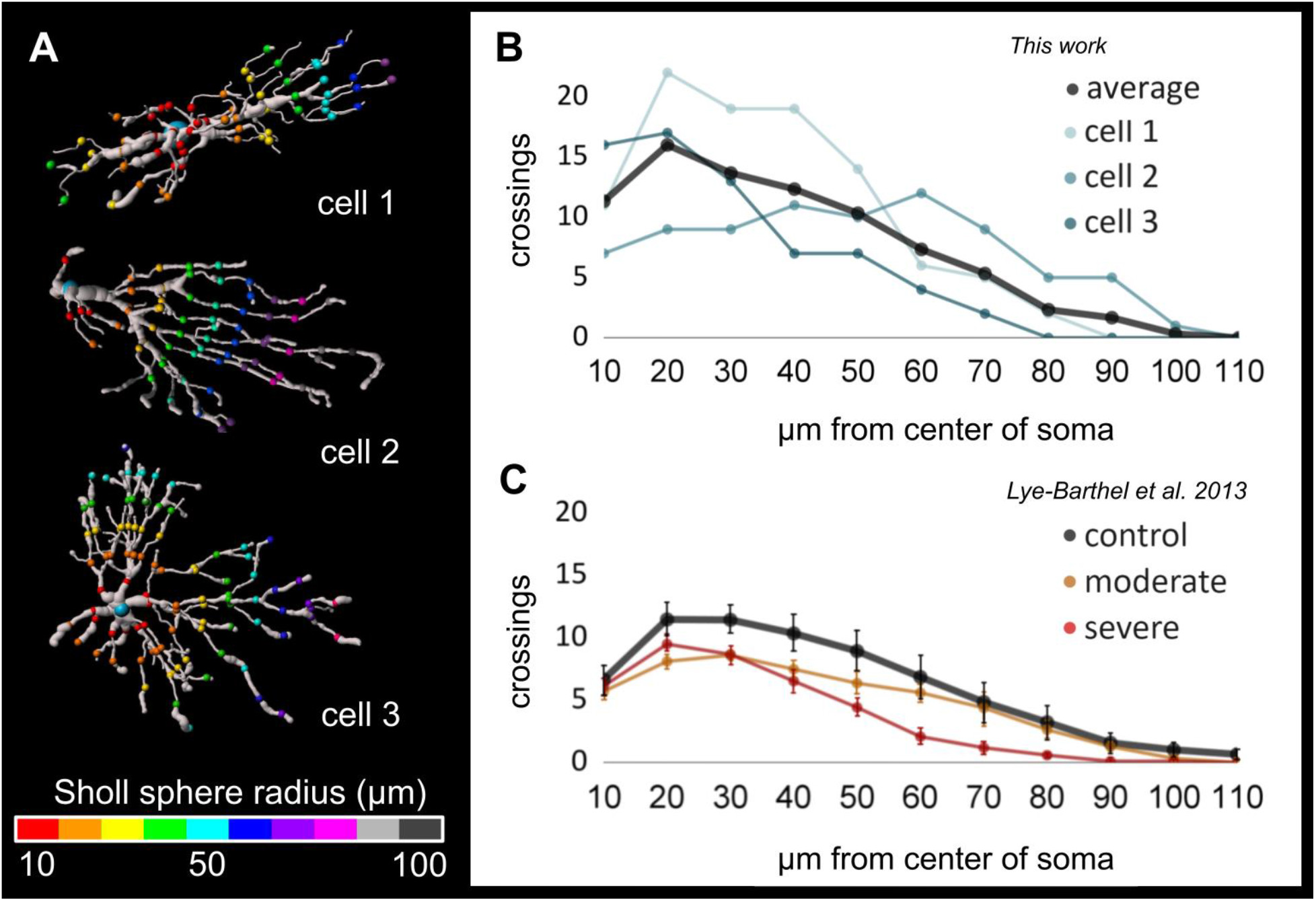
Astrocyte models allow 3D Sholl analysis. **A)** Three representative goat astrocytes were analyzed in 3D. Colored points indicate branch crossings with Sholl spheres created at 10 μm increments from the astrocyte soma center-point. **B)** Crossing counts of the three example cells and the average. **C)** Astrocyte branch crossings at Sholl spheres in mouse eyes from Lye-Barthel et al. 2013.(Lye-Barthel et al., 2013) The crossing counts were higher in the collagenous LC, although the pattern of distribution was remarkably consistent with that of the control mouse.

The morphologies of collagenous LC astrocytes were in line with those of other fibrous astrocytes from large mammals, just as glial lamina astrocyte morphologies are in line with fibrous astrocyte morphologies of the mouse.(Oberheim et al., 2009; Sun et al., 2010) Protoplasmic astrocytes found in other areas of the central nervous system have substantially bushier morphologies and higher branch densities than do fibrous astrocytes.(Oberheim et al., 2009)

### 3.5. Labeling and visualization of LC astrocytes is effective across species

MuDi allowed visualization of LC astrocytes across all species with a collagenous LC tested (**Figure 8**). Astrocytes were visualized in the LC of monkey (**Figure 8A**), goat (**Figure 8B**), pig (**Figure 8C**), and sheep (**Figure 8D**). Detail of astrocyte morphologies was visible across species, including thin higher-order branches and end-feet (A’-H’). End-feet were observed in contact with collagenous LC beams (**Figure 4B, Figure 8**).

**Figure 8.**
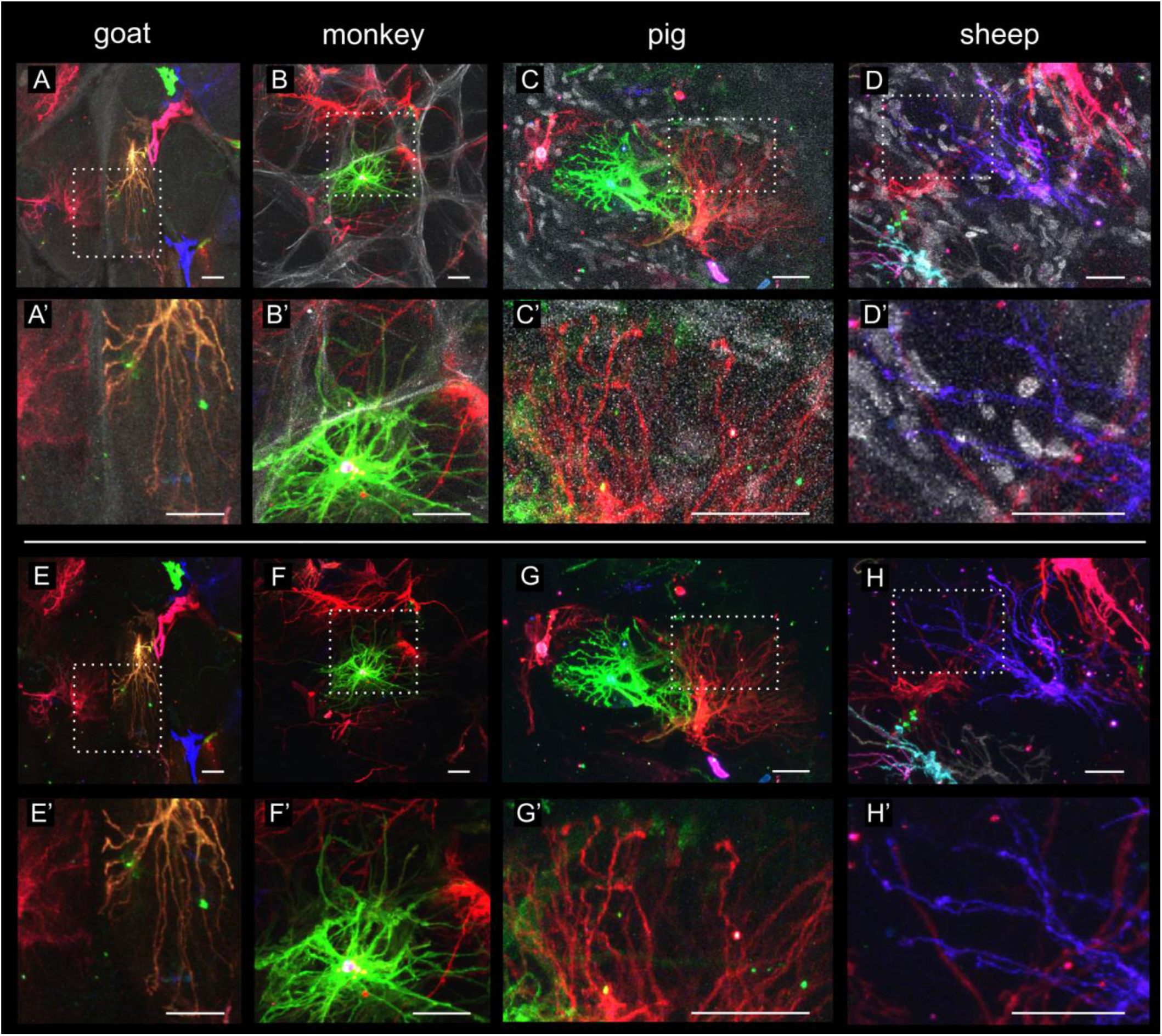
Labeling and visualization of LC astrocytes are effective across species. LC astrocytes were successfully MuDi labeled and visualized within the goat **(A, E)**, monkey **(B, F)**, pig **(C, G)**, and sheep **(D, H)** LC. Representative example images demonstrate labeling of the soma, fine processes, and astrocyte end-feet. Detail within the white boxes of **A-D and E-H** is shown in **A’- D’ and E’-H’**. Sheep and pig samples in these images were counterstained with DAPI. The grayscale channel in goat and monkey images show tissue autofluorescence only. Scale bars: 25 μm.

### 3.6. Multicolor DiOlistics is compatible with labeling and imaging of other structures

MuDi was compatible with labeling and imaging of other collagenous LC structures. We have shown that MuDi in the LC allows for seven color channels with which astrocytes can be visualized. Paired SHG, DAPI labeling, and additional post-DiOlistics markers, this provides the potential for 12 distinct channels for visualization of LC structures in the same tissue.

SHG imaging and visualization of collagen was possible in DiOlistically-labeled LC tissues without additional sample labeling or preparation. The presence of LC collagen beams is a defining feature of the anatomic region and is a key difference between the collagenous LC, found in humans and the species used in this study, and the glial lamina of mice and rats. Visualizing LC collagen in tandem with astrocyte morphology and spatial arrangement can provide verification of tissue location and a better understanding of individual astrocyte relationships to surrounding collagen beams.

With many fluorophores available being in the DiO, DiI, and DiD ranges (similar to FITC, TRITC, and Cy5), the use of MuDi could preclude the use of other common labeling options. However, due to the solubility of DiO, DiI, and DiD in ethanol, these dyes could be washed from tissue samples in an ethanol gradient. Samples labeled for GFAP (FITC-range secondary) and lectin (Cy5-range conjugate) that were then washed of DiOlistics dyes allowed specific visualization of GFAP intermediate filaments and blood vessels without appreciable DiOlistics signal (**Figure 9**). This provided an additional two distinct channels for visualization of LC structures in these tissues, with potential for an additional third post-DiOlistics channel in the TRITC/DiI-range, not used here. A total of 12 distinct channels for visualization of LC structures could be possible in the same sample with the use of MuDi (7), DAPI (1), SHG (1), and additional labels after washing away DiOlistics dyes in the FITC, TRITC, Cy5 ranges (3).

**Figure 9.**
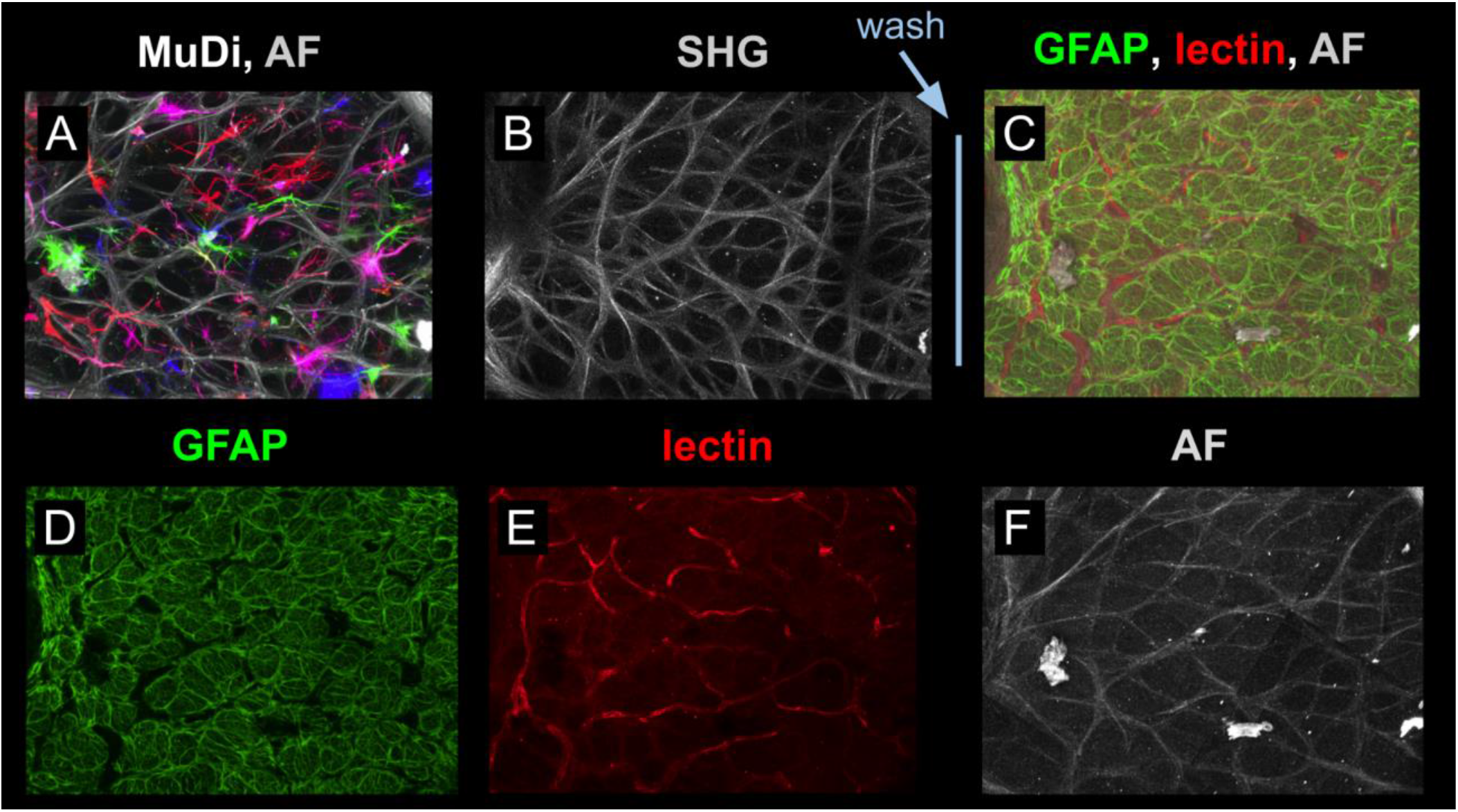
DiOlistic labeling is compatible with downstream labeling and imaging of other structures. **A**) A slice through the monkey ONH containing collagenous LC labeled with MuDi. The gray 405nm channel shows tissue autofluorescence. **B**) The same sample was imaged via second harmonic generation (SHG) microscopy to visualize collagen. **C**) Cell membrane dyes were washed from the sample with ethanol prior to re-imaging. The sample was labeled with GFAP and tomato lectin to visualize astrocyte intermediate filaments and blood vessels. Signal from GFAP, lectin, and tissue autofluorescence are shown independently in **D, E**, and **F**, respectively. LUTs in **E** and **F** were adjusted to aid visualization. Example tissues were of monkey origin.

### 3.7. Multicolor DiOlistic labeling can reveal axons and other cell types

MuDi preferentially labels astrocytes in LC coronal slices due to their abundance, size, and general orientation along the coronal plane. However, this labeling technique is not inherently discriminatory of cell type. RGC axons, LC cells, and scleral fibroblasts can also be labeled by this technique but have markedly different labeling frequencies and cell morphologies. Examples of labeled cells with fibroblast-like morphologies along LC beams and in the peripapillary sclera region are included in **Figure 10**. Because of the largely longitudinal orientation of RGC axons, individual axons will occupy little space in the coronal plane. When axons in coronal slices are labeled with DiOlistics, they appear as small puncta or hollow tubes in projection, depending upon magnification power. With increasing deviation of sample orientation from the coronal plane, each axon occupies more space within a tissue slice. Axons are more likely to be stochastically hit by a DiOlistics microcarrier when they occupy more space. Axon labeling in slices cut at an angle that deviated from the coronal plane is shown in **Figure 11**. Axons are visualized in the nerve fiber layer and prelamina (**Figure 11A**) as well as the LC, retrolamina, and optic nerve (**Figure 11B, C**). In samples sectioned longitudinally, axons were frequently labeled, apparent as filamentous structures oriented along the sagittal plane (**Figure 11C**).

**Figure 10:**
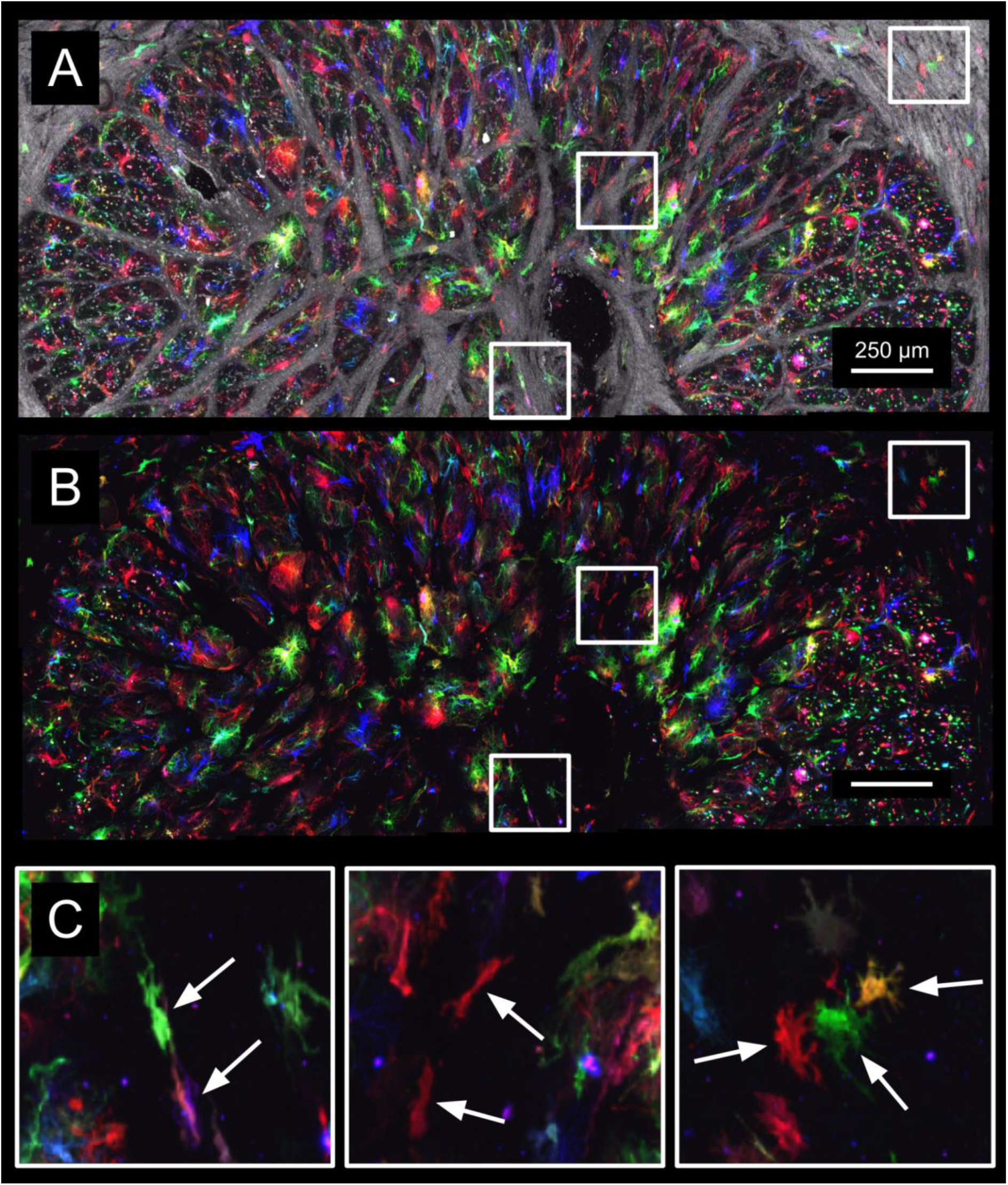
Multicolor DiOlistic labeling can reveal cells with fibroblast-like morphologies. **A)** MuDi-labeled sheep LC with extracellular matrix autofluorescence shown in grayscale. **B)** The same image as above, showing MuDi signal only. White boxes within **A** and **B** indicate regions containing presumptive fibroblasts along collagen beams and in the peripapillary sclera. **C)** Detail of MuDi-labeled cells with fibroblast-like morphologies (arrows).

**Figure 11.**
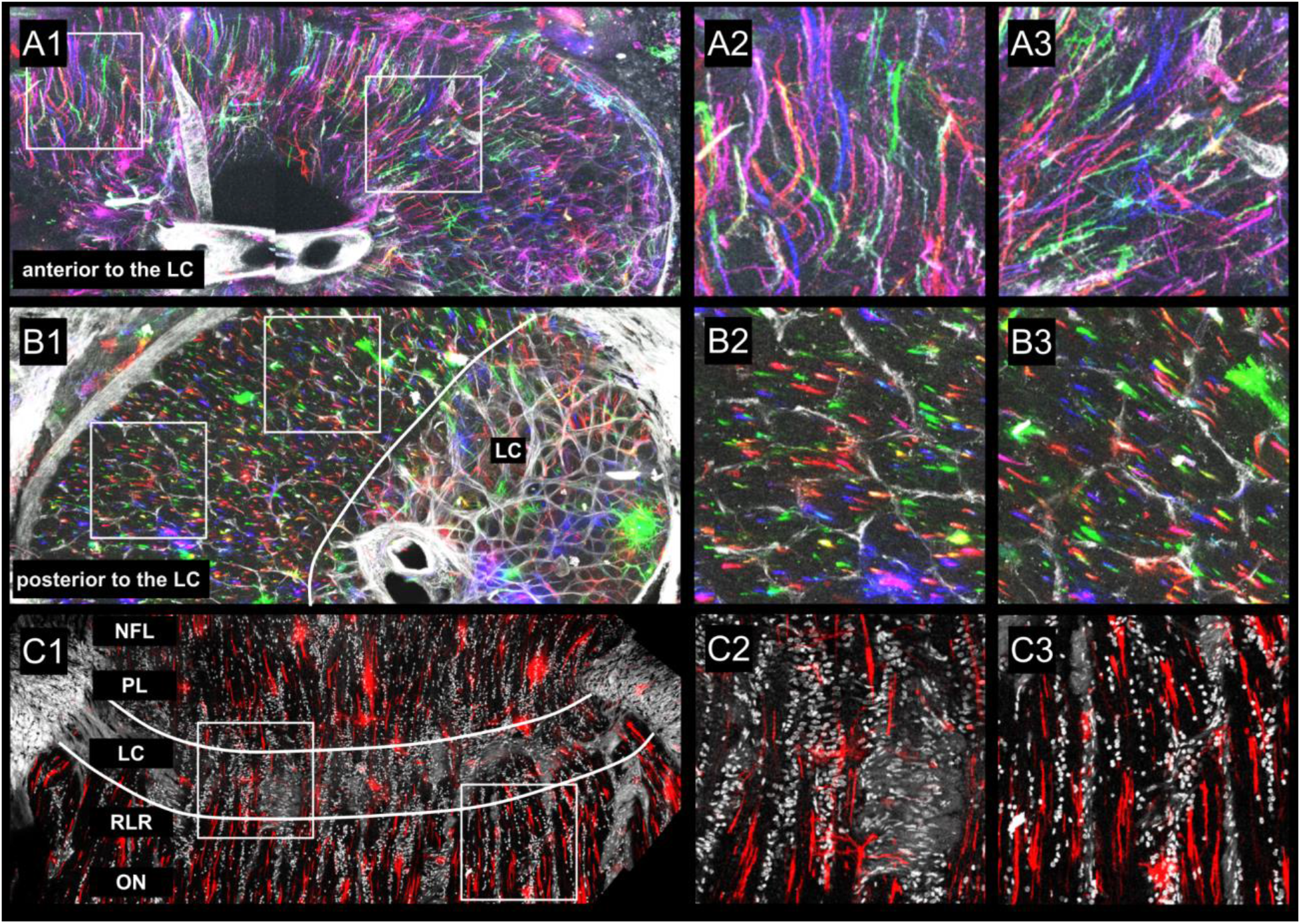
Multicolor DiOlistic labeling can reveal axons DiOlistic labeling can reveal axons. In tissues anterior **(A)** and posterior **(B)** to the LC, DiOlistically-labeled RGC axons can be seen running largely parallel to each other. Axons are most readily labeled and visualized in longitudinal sections **(C)**, where single axons occupy the greatest amount of area, increasing the probability of DiOlistic axon labeling and dye diffusion across greater length of the axon. Samples in **A** and **B** were from coronal slices of monkey, labeled with MuDi. Grayscale in **A** and **B** shows tissue autofluorescence from the 405nm channel only. The longitudinal slice in **C** was from goat, labeled with single-color DiI DiOlistics and counterstained in DAPI. Although axons are more readily visualized in these samples than in coronal slices through the LC, not all structures labeled via DiOlistics here are axons. For example, the coronally-oriented DiI-labeled branches in **C2** may belong to LC astrocytes. NFL: nerve fiber layer, PL: prelamina, LC: lamina cribrosa, RLR: retrolaminar region, ON: optic nerve.

## 4. Discussion

In this work, we investigated the capability of MuDi labeling to allow visualization of astrocytes in the collagenous LC. Analysis of tissues treated with this approach resulted in three key findings. First, DiOlistics allowed visualization of individual astrocytes and fine details of their morphologies across the collagenous LC. Second, multicolor labeling granted visualization of multiple cells in close proximity at once while still allowing to discern individual cells from their neighbors. Third, 3D astrocyte models made from confocal image volumes facilitated automated quantification of morphological features. Below we discuss the significance and implications of these findings.

### 4.1. DiOlistics allowed visualization of individual astrocytes and fine details of their morphologies across the collagenous LC

As shown in earlier works from other groups, DiOlistic labeling is capable of revealing complex morphologies of neurons in brain tissues(O’Brien and Lummis, 2006; Seabold et al., 2010; Staffend and Meisel, 2011) down to the detail of their dendritic spines. DiOlistics is also capable of labeling astrocytes in complex tissue environments, as shown in the mouse, chimpanzee, and human brains.(Oberheim et al., 2009, 2008) In mice, DiOlistics has been used to reveal individual astrocyte morphologies in the glial lamina specifically.(Gan et al., 2000; Oberheim et al., 2008) Individual lamina astrocyte morphologies in species with a collagenous LC, like humans, are less well-described. We applied DiOlistics to the collagenous LC of pig, sheep, goat, and monkey to visualize resident astrocyte morphologies. We demonstrated that DiOlistics allowed visualization of individual astrocytes and microscale detail of their morphologies in the context of their complex LC tissue environments.

As shown in **Figure 4A**, DiOlistic labeling resulted in transfer of dye-coated microcarriers to cells across coronal slices through the LC. The same sample imaged at higher magnification (**Figure 4B**) reveals morphologies of individual astrocytes within LC pores. In samples immunostained for GFAP, a common astrocyte marker, DiOlistically-labeled cells with astrocyte morphologies were GFAP^+^. Detailed structural analysis of GFAP, F-actin, nuclei, and collagen beams in the healthy and glaucomatous human LC(Guan et al., 2022) has provided important insights into the differences of these elements between health and advanced disease. However, detail of individual LC astrocyte morphologies cannot be determined through visualization of these elements alone. Consistent with other descriptions of ONH astrocytes observed with different methods,(Ju et al., 2015; Lye-Barthel et al., 2013; Nguyen et al., 2017; Wang et al., 2017) the LC astrocytes visualized in this study had multiple ramified processes. Astrocyte end-feet were often observed in contact with LC beams, shown in detail in **Figure 7**. To our knowledge, individual astrocyte morphologies in the collagenous LC, labeled through a method that can reveal the entirety of the cell plasmalemma, have not been previously reported.

Primary culture of astrocytes from models that have a collagenous LC has been accomplished(Hernandez et al., 1988; Lopez et al., 2020) and provides an excellent opportunity for *in vitro* visualization. However, the process of primary culture requires removal of cells from the context of surrounding collagen beams, retinal ganglion cells, blood vessels, and other astrocytes. This, along with maintenance on a 2D surface rather than a 3D tissue environment, can result in substantial alteration of astrocyte morphologies. Visualization of LC astrocytes within the context of their environment, as we have done here, with preserved spatial relationships to surrounding structures, is optimal to understand their original morphologies.

### 4.2. The multicolor approach allows for labeling of many cells at once while still allowing separation of cells from their neighbors

Multicolor DiOlistics, developed by Gan et. al, was demonstrated to allow visualization of and spatial separation of neighboring neurons and astrocytes in the mouse brain and retina.(Gan et al., 2000) We applied this multicolor approach to label and visualize tens to hundreds of cells across the pig, goat, sheep, and monkey LC at once while allowing separation of individual cells from their neighbors. Due to the density of astrocytes in the LC and their complex, highly branched morphologies, techniques using a single fluorescent label to visualize astrocytes would require a relatively low labeling density in order to effectively distinguish between neighboring cells. Manual injection of different dyes into individual cells is possible,(Ju et al., 2015) but can be time and labor-intensive. After a single gene gun shot, MuDi allows spatial separation of neighboring astrocytes to be visualized through multiple distinct color channels.

MuDi is similar in concept to Brainbow small research animal reporter lines(Livet et al., 2007; Pan et al., 2011; Richier and Salecker, 2015) which have been used to help distinguish individual neurons and astrocytes from their neighbors in the brain. As in the *Thy1-Brainbow-1*.*0* mouse line, a greater number of color channels than seven can be created from 3 fluorescent reporters if more intermediate colors are utilized (ex. two parts green and one part red). With combinatorial expression of three fluorescent reporters (GFP, RFP, and CFP), Livet et al could distinguish neural cells in approximately 100 distinct color groups.(Livet et al., 2007) MuDi, and variations of it, can be useful to investigate questions similar to those allowed by Brainbow tissues, but in large animal or human donor eyes where gene modification is not as easily achieved. For example, the spatial territories of individual collagenous LC astrocytes and the corresponding regions of RGC axons over which they have direct physiologic influence could be observed in health in disease. This has potential to inform our understanding of how individual astrocyte territories and their zones of physiologic influence may change in stress states, informing our understanding of disease progression and pathways for neuroprotective strategies.

### 4.3. Automated quantification of morphological features was facilitated by DiOlistics-derived 3D astrocyte models

3D astrocyte models were readily constructed from confocal image volumes for automated quantification of morphological features. Sharing representative images and qualitative interpretations based on large image sets can be a valuable way to communicate summaries of scientific findings. However, quantifying morphological features from large image datasets in addition to sharing qualitative interpretations can allow for a clearer, broader, and more replicable picture of LC astrocyte morphology. Additionally, morphologies of astrocytes are often categorized into two broad, qualitative groups: healthy / “non-reactive” and pathologic / “reactive.” As the understanding of astrocyte structure and function has become more nuanced, it is important to supplement qualitative and categorical observations with detailed quantitative ones that can be evaluated and analyzed statistically, as is allowed by the approach described in this work.

The quantification of morphological features that this approach allows opens doors to ask quantitative questions about the heterogeneity of astrocyte morphologies within a single individual, among different individuals, and between different species, as we aim to do in our future work. The technique can allow for investigation of quantitative differences and similarities in astrocyte morphologies between LC regions, such as central vs. peripheral LC astrocytes or nasal vs. temporal ones. It can additionally allow for quantification of astrocyte spatial relationships with surrounding collagenous LC elements, such as collagen beams, blood vessels, and other astrocytes. Importantly, it may provide clearer insights into astrocyte morphology throughout the progression of aging and glaucoma in experimental models or in human donor eyes. The ability to label tens to hundreds of cells at once in a single tissue and quantify their morphologies can allow us to discern patterns that may be missed in smaller sample sizes or with qualitative interpretation alone.

Important limitations of this approach must be acknowledged. DiOlistics relies on stochastic labeling of cell membranes and is not inherently discriminant of cell type. Astrocytes are abundant in the LC, accounting for over 90% of nuclei in the region,(Guan et al., 2022) with cell bodies largely oriented along the coronal plane. Therefore, many cells labeled in these coronal slices were astrocytes. MuDi will label larger cells or more coronally-oriented cells more frequently than smaller and more longitudinally-oriented ones. This approach may therefore oversample larger astrocytes oriented along the coronal plane and undersample smaller astrocytes with a smaller footprint along the coronal plane. Despite the lack of complete cell-type specificity, astrocyte morphologies were distinctly discernible from those of other LC cellular elements, such as retinal ganglion cell axons and LC cells. The technique’s compatibility with other more cell-type-specific labeling methods, such as the GFAP labeling in this study, allowed for additional validation of the identity of DiOlistically-labeled cells.

The penetration of microcarriers in tissues is determined by the biomechanical properties of the tissue, the pressure at which microcarriers are propelled into the tissue, and the distance the gene gun is positioned from the sample. At 150-200 PSI and gene gun distance of 30mm, we observed microcarriers up to 80μm deep within the samples. Higher pressures can increase microcarrier penetration depth, but risks damage to tissues. The limited penetration of microcarriers necessitates sectioning of tissues prior to labeling. The process of sectioning can result in severing cells and incomplete visualization of cell morphology. The collection of thick tissue slabs with a vibratome minimizes the number of cells in samples with severed processes. However, severed cells are likely to leak dye into surrounding tissues more quickly than intact cells. Cells with poorly defined morphologies due to dye leakage were not suitable for 3D model construction and can be excluded from morphological analyses. Gene gun modifications have been shown to allow for deeper bullet penetration at lower gas pressures.(O’Brien et al., 2001) A combination of gene gun modification and dye-preserving tissue clearing methods(Ke et al., 2016; Li et al., 2019) could allow for visualization of labeled astrocytes in whole LCs.

The time-sensitive nature of imaging samples labeled with MuDi is an important consideration for study design in which large high-resolution confocal image sets can take several hours to collect. LC slices labeled with MuDi must be imaged within 24-48 hours of labeling, as dyes can leak out of cells and into surrounding space over time. Dye leakage and redistribution obscures the morphology of individual cells, preventing morphological analysis. Examples of tissues with dye leakage preventing high quality visualization of individual cell morphologies are included in **Supplemental Figure 2**. The use of fixable dyes such as CM-DiI may help mitigate this issue.

The tissues used in this study were fixed prior to labeling and imaging. This prevents investigation of cell function in conjunction with morphology and tracking any changes over time. DiOlistic labeling has been accomplished in organotypic slice cultures of brain tissues. Preliminary data from our lab demonstrates that MuDi labeling can be accomplished in the non-fixed LC vibratome slices as well. This technique, or similar techniques using fluorescent dextrans instead of cell membrane dyes,(Jakobs, 2014) may allow investigation of LC astrocyte structure and function in *ex vivo* slice culture. *Ex vivo* culture and drug delivery systems of eyes with a collagenous LC(Chan et al., 2022) combined with DiOlistics may be able to relate structural astrocyte features to functional ones and track changes over time.

Overall, MuDi labeling of the collagenous LC provides a way to visualize morphological detail of individual LC astrocytes at a scale not provided by other techniques. This approach revealed LC astrocyte morphologies, allowed cells in close proximity to be distinguished from one another, and facilitated quantification of morphological features. It is compatible with other labeling and imaging techniques such as immunohistochemistry and SHG. MuDi labeling provides the potential for future large-scale qualitative and quantitative analysis of astrocyte morphological features and spatial relationships with interacting elements, such other astrocytes, collagen beams, blood vessels, and axons, across the collagenous LC. Investigation was possible across different healthy animal models. In future studies, this technique may allow for investigation of LC astrocyte morphology in different glaucoma model progression states or in human donor tissues as well. This approach can help provide important insights into collagenous LC astrocyte physiology in health and disease.

## Acknowledgements

Dr. Martin Oberbarnscheidt, for generously sharing vibratome equipment and training in its proper use

## Funding

Supported in part by National Institutes of Health grants R01-EY023966, R01-EY031708, R01-HD045590, R01-HD083383, 1S10RR028478-01, P30-EY008098, and T32-EY017271; Eye and Ear Foundation (Pittsburgh, PA); Research to Prevent Blindness (unrestricted grant to UPMC Ophthalmology and Stein Innovation Award to Sigal IA).

We would like to acknowledge Dr. Martin Oberbarnscheidt for generously sharing vibratome equipment and training in its proper use.

## Supplemental Materials

**Supplemental Figure 1.**
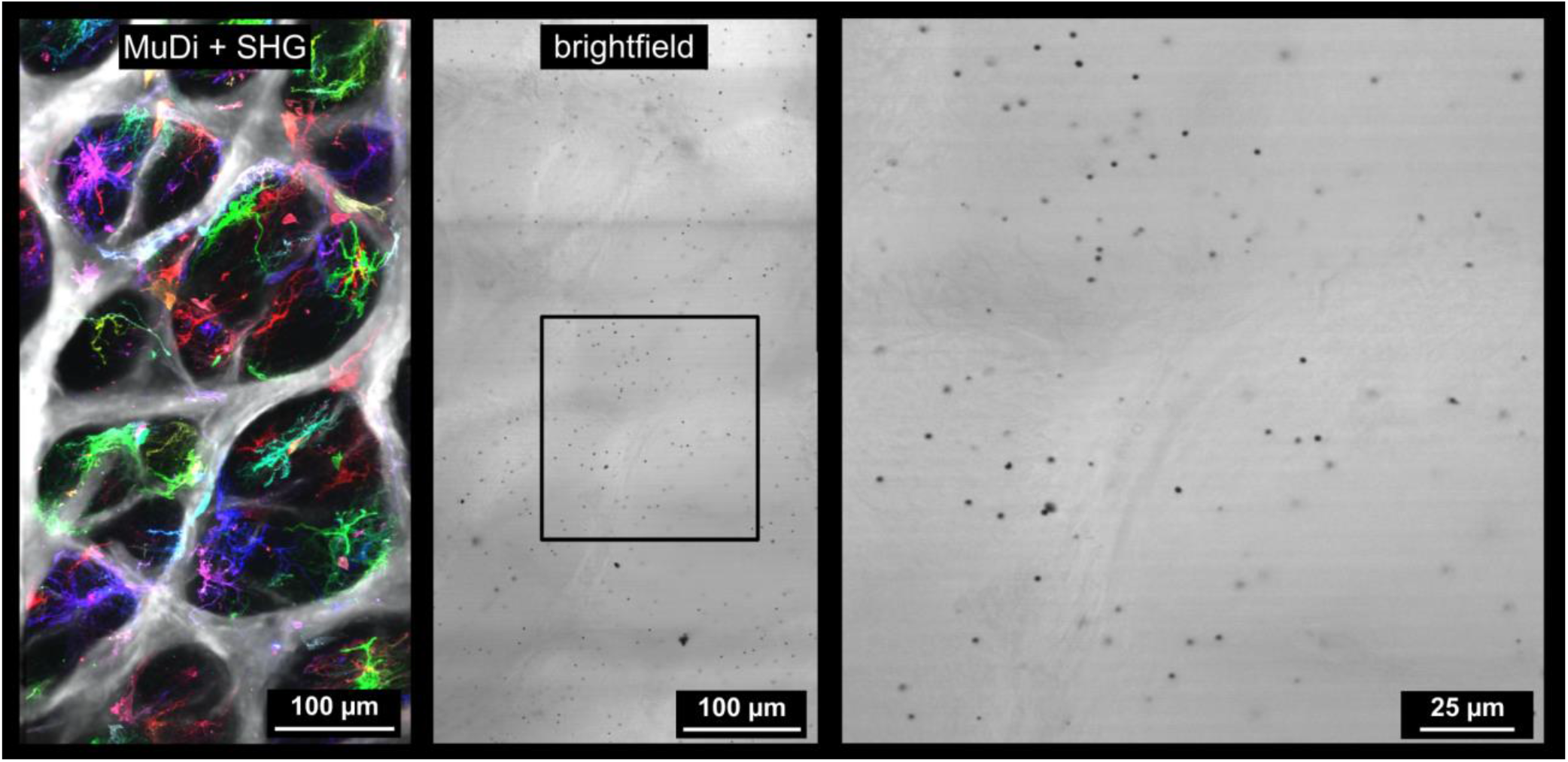
Visualization of microcarrier distribution. An image of the region shown in **Figure 4B** (left) is provided as a reference for the respective brightfield image (middle). The brightfield image shows boundaries of the same collagen beams and neural tissue pores. Detail of the region within the black box in the middle panel shows bullets from multicolor DiOlistics visible as dark puncta approximately 1μm in diameter. Optimal labeling density for revealing a large number of cells while minimizing spatial overlap among those in the same color channel was approximately 140 cells per mm^2^ of canal section area. SHG = second harmonic generation.

**Supplemental Figure 2:**
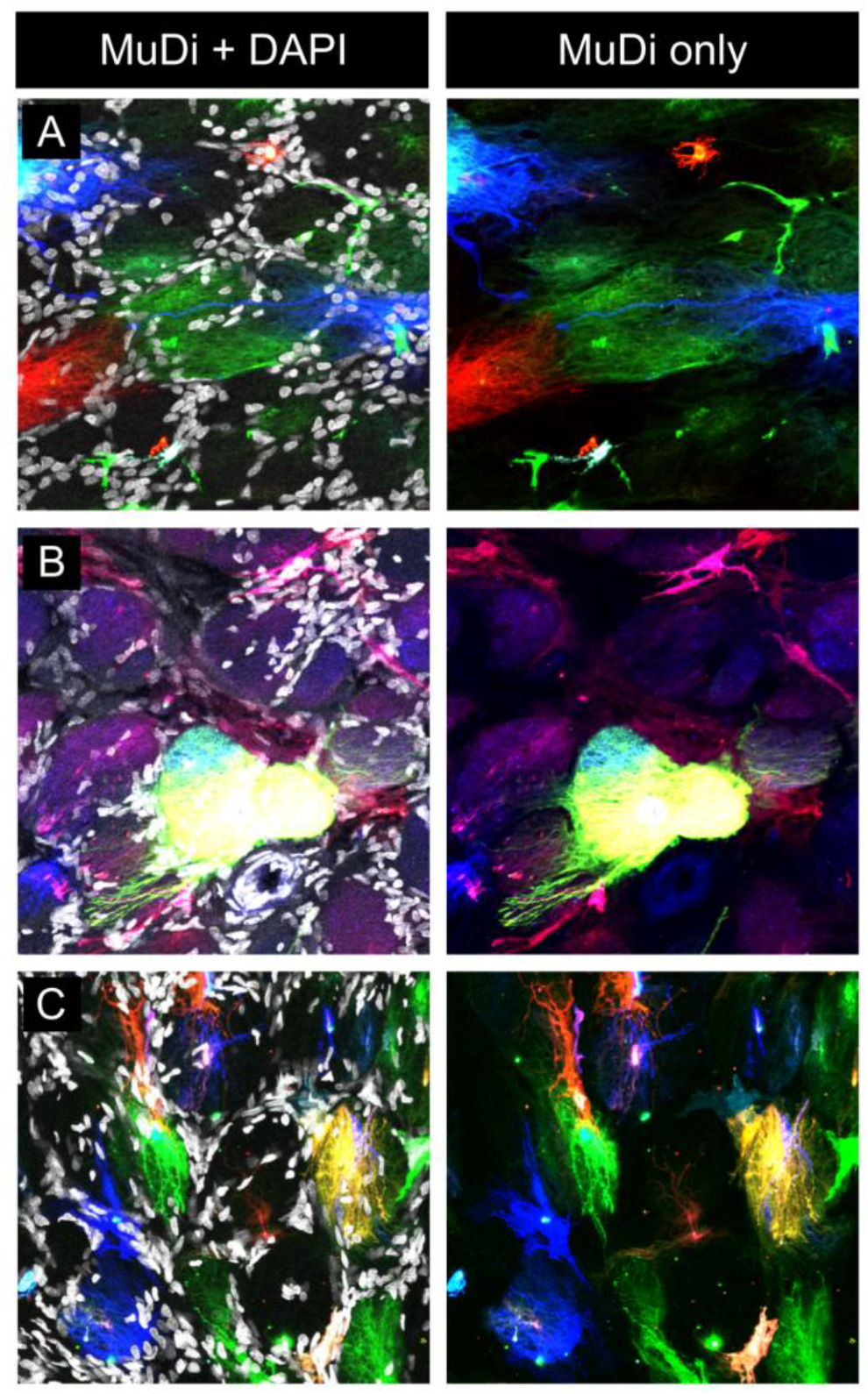
Dye leakage obscures visualization of morphology. Example images include tissues labeled with MuDi in which individual astrocyte morphologies were not distinguishable. Dyes can diffuse from individual labeled cells into surrounding areas if cell membrane integrity is sufficiently compromised. Diffuse dye-labeled regions with multiple nuclei and without clear astrocyte morphologies, apparent in **A**, were not considered astrocytes and were excluded from morphological analysis. Microcarriers can be aberrantly delivered in clumps if not properly sonicated, filtered, or stored with desiccant. Delivery of microcarrier clumps can result in large patches of tissue being labeled with the same dye (such as the green region in **B**) and potential lysis of cells bombarded with microcarrier clumps larger than 1μm in diameter. Some dye diffusion was confined to individual pores, as in **C**, labeling parts of several cells within a pore. Other dye spread, particularly in samples treated with fixative that contained methanol or labeled at pressures higher than 200 PSI, was more diffuse across the sample, such as the blue and red signal in **B**.

## Videos

**Video 1**:

Example 3D DiOlistics confocal volume information shown in relation to a 3D astrocyte model.

**Video 2**:

Astrocyte models from **Figures 5**-**7** shown in 3D.

