## Supplemental Materials for "Individual Astrocyte Morphology in the Collagenous Lamina Cribrosa Revealed by Multicolor DiOlistic Labeling"

**Short title:** MuDi labeling individual collagenous LC astrocytes

* **Correspondence:**

Ian A. Sigal, Ph.D.
Laboratory of Ocular Biomechanics
Department of Ophthalmology, University of Pittsburgh School of Medicine
203 Lothrop Street, Eye and Ear Institute, Rm. 930, Pittsburgh, PA 15213

www.OcularBiomechanics.com

**Keywords**: astrocyte, lamina cribrosa, microscopy, morphology, optic nerve head

**Disclosures:** Nothing to disclose.

**Funding:** Supported in part by National Institutes of Health grants R01-EY023966, R01-EY031708, R01-HD045590, R01-HD083383, 1S10RR028478-01, P30-EY008098, and T32-EY017271; Eye and Ear Foundation (Pittsburgh, PA); Research to Prevent Blindness (unrestricted grant to UPMC Ophthalmology and Stein Innovation Award to Sigal IA).

**Acknowledgements:** Dr. Martin Oberbarnscheidt, for generously sharing vibratome equipment and training in its proper use

**Supplemental Figure 1**) Visualization of microcarrier distribution

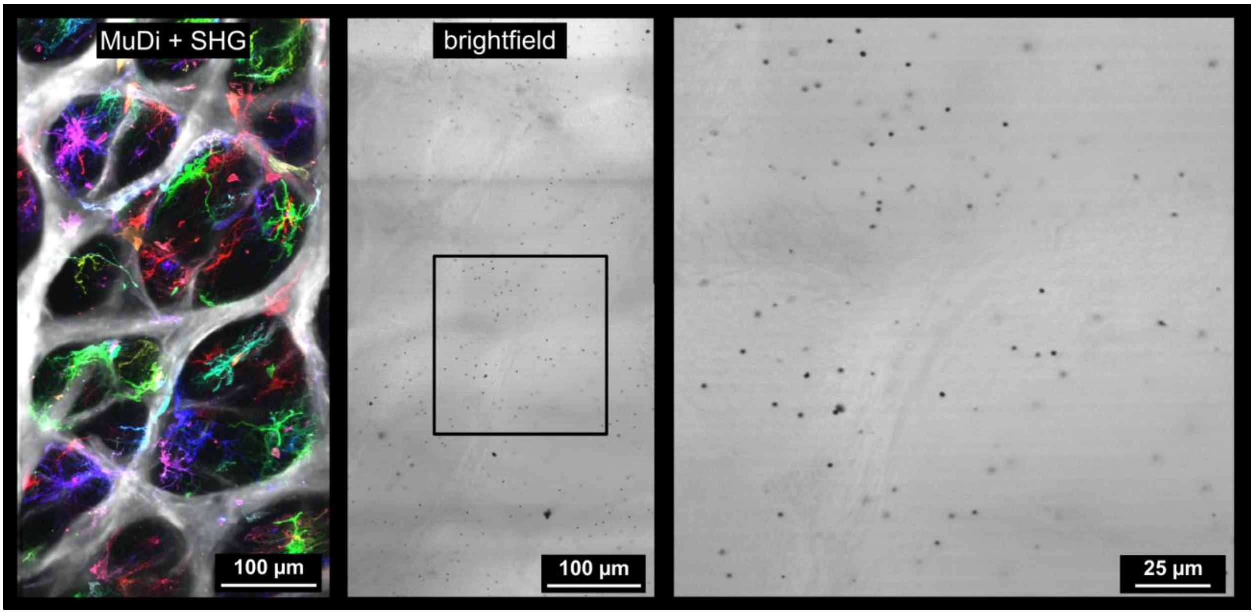


**Supplemental Figure 1**) Visualization of microcarrier distribution. An image of the region shown in **Figure 4B** (left) is provided as a reference for the respective brightfield image (middle). The brightfield image shows boundaries of the same collagen beams and neural tissue pores. Detail of the region within the black box in the middle panel shows bullets from multicolor DiOlistics visible as dark puncta approximately 1μm in diameter. Optimal labeling density for revealing a large number of cells while minimizing spatial overlap among those in the same color channel was approximately 140 cells per $mm^2$ of canal section area. SHG = second harmonic generation.

**Supplemental Figure 2**) Dye leakage obscures visualization of morphology.

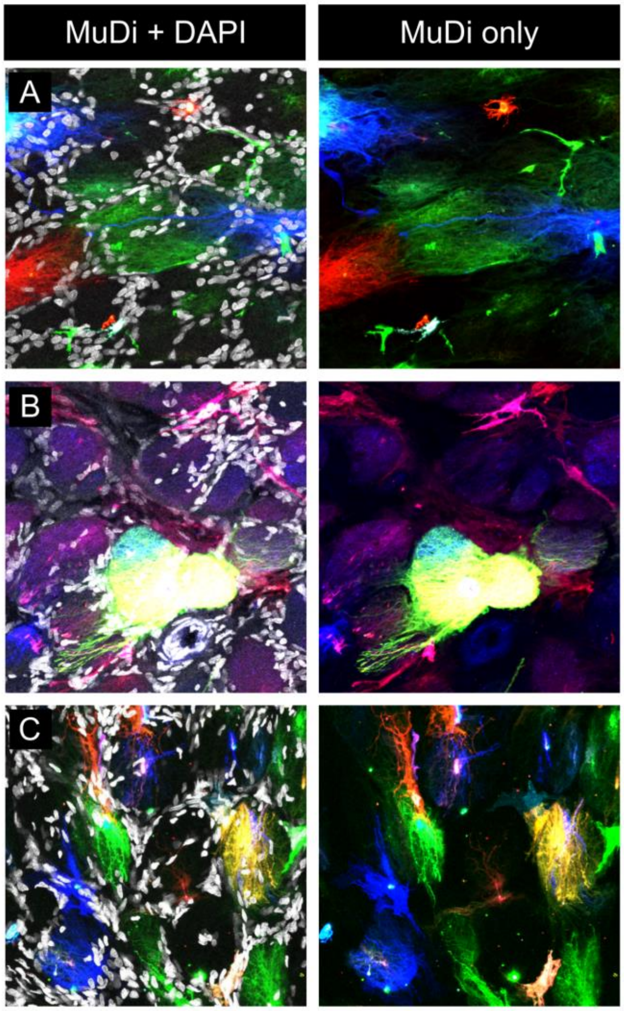


**Supplemental Figure 2**: Dye leakage obscures visualization of morphology. Example images include tissues labeled with MuDi in which individual astrocyte morphologies were not distinguishable. Dyes can diffuse from individual labeled cells into surrounding areas if cell membrane integrity is sufficiently compromised. Diffuse dye-labeled regions with multiple nuclei and without clear astrocyte morphologies, apparent in **A**, were not considered astrocytes and were excluded from morphological analysis. Microcarriers can be aberrantly delivered in clumps if not properly sonicated, filtered, or stored with desiccant. Delivery of microcarrier clumps can result in large patches of tissue being labeled with the same dye (such as the green region in **B**) and potential lysis of cells bombarded with microcarrier clumps larger than 1µm in diameter. Some dye diffusion was confined to individual pores, as in **C**, labeling parts of several cells within a pore. Other dye spread, particularly in samples treated with fixative that contained methanol or labeled at pressures higher than 200 PSI, was more diffuse across the sample, such as the blue and red signal in **B**.

**Videos:**

**Video 1**:
Example 3D DiOlistics confocal volume information shown in relation to a 3D astrocyte model.

**Video 2**:
Astrocyte models from **Figures 5-7** shown in 3D.
